# High frequency soil polarization can phenotype crop roots noninvasively

**DOI:** 10.1101/2023.01.12.523853

**Authors:** Huijie Gu, Imre Cseresnyés, John R. Butnor, Baoru Li, Benjamin Mary, Hongyong Sun, Xiying Zhang, Yang Lu, Xiuwei Liu

## Abstract

Noninvasive and nondestructive root phenotyping techniques under field conditions are sorely needed to advance plant root science. Soil polarization measured by electrical capacitance (EC_soil_) has the potential to meet this requirement, but whether it specifically detects root properties remains unexplored. We carried out manipulative experiments where wheat (*Triticum aestivum* L.) and maize (*Zea mays* L.) roots were buried in soil or immersed in hydroponic solution combined with pot trials to reveal the mechanism of root trait detection by EC_soil_, while a field experiment was conducted to test its feasibility to determine root depth distribution. We found that EC_soil_ measured at low current frequency (< 1 kHz) was not significantly affected by the addition of roots to the system either by burying roots in soil or immersing them in solution. At frequency greater than10 kHz a shift occurred, and root polarization contributed more to EC_soil_ which was positively correlated with root volume. When EC_soil_ was measured at high frequency (30 kHz −100 kHz) it was well correlated with root volume vertical distribution in the field. The measurement error after soil moisture calibration at depths of 10 cm, 20 cm, 30 cm and 40 cm was 0.4%, 12.0%, 1% and 34%, respectively. Our results demonstrate that EC_soil_ is a robust method to measure *in situ* root distribution and we believe the newly available high frequency measurement equipment combined with novel root prediction models will enable EC_soil_ to be widely used for root phenotyping in the future.

## 1. Introduction

Roots are responsible for anchoring plants, nutrient, and water uptake, as well as soil carbon and nitrogen cycling via root exudates and fine root turnover (Lynch, 2019; Karlova *et al*., 2021). Improving the utilization efficiency of crop water and fertilizer is important for increasing crop production and can be augmented by selecting genotypes with desirable root traits for given environment (Lynch, 2013; Lynch, 2019; Karlova *et al*., 2021). However, the lack of in-situ nondestructive, noninvasive, and high-throughput crop root phenotyping methods prevents the optimization of root function.

Electrical methods including electrical capacitance (EC), electrical resistance, electrical impedance tomography, electrical resistivity tomography and spectral induced polarization approaches etc. are noninvasive, nondestructive, and can be deployed rapidly for high-throughput applications (Mancuso, 2012; Cimpoiasu *et al*., 2020; Ehosioke *et al*., 2020). EC is the amount of charge stored in a capacitor under a certain voltage. Generally, the electrical methods such as EC can be divided into stem-based (root EC (EC_root_), one electrode is located on the stem and the other one is in the medium) measurement and medium-based (medium capacitance, both electrodes are located in the medium) measurement (Tsukanov and Schwartz, 2021).

EC_root_ predicts root properties such as root surface area, root dry mass and root volume by measuring the EC between stem and substrate (Chloupek, 1972; Dalton, 1995; Ehosioke *et al*., 2020). It is hypothesized that the electrical current flows from the stem into the root system, exiting the root hair zone and continuing to the soil media (Dalton, 1995). Therefore, the root system can be regarded as a capacitor, and the measured capacitance is proportional to the polarized membrane (dielectrics) surface area. After Chloupek (1972) found that EC_root_ was closely related to root traits, it was widely used to estimate root traits. However, electrical current can escape from the surface of roots calling ability of EC_root_ to quantify root size into question (Peruzzo *et al*., 2020; Urban *et al*., 2011). Gu *et al*. (2021) further demonstrated that EC_root_ can estimate root traits best under dry soil conditions.

When the medium containing roots receives an low frequency (< 10 kHz) alternating current (AC), counterion polarization of root cells due to the existence of electrical double layer would increase the dielectric constant of the medium (Weigand and Kemna, 2017; Weigand and Kemna, 2019; Tsukanov and Schwartz, 2021). As a result of this phenomenon, root characteristics may be sensed by medium EC. Mary *et al*. (2017) found that burying coarse root samples (diameter > 2 mm) in the soil would increase soil polarization response, and the amount of additional root volume was directly related to increased soil polarization. Furthermore, Tsukanov and Schwartz (2020) found that the polarization signal of nutrient solution was positively correlated with root dry mass and root surface area and proving that immersed roots were the main source of polarization in hydroponic systems. Liao *et al*. (2015) used EC tomography at 1 kHz to successfully image the position of *Rohdea japonica* roots in hydroponic solution. These studies indicated that root size and its distribution can be estimated by the medium polarization. However, the measuring frequency in these previous studies was relatively low (<10 kHz). Bulk soil polarization is composed of root polarization and soil colloid polarization (Kessouri *et al*., 2019), the volume fraction of crop roots in soil is low (for example, around 0.006 for wheat, 0.02 for maize, 0.006 for rice etc. in topsoil) meaning the root polarization tends to be minimal (Gan *et al*., 2011) at low frequency. Plant root growth and root uptake affect the soil water, ion content and porosity which in turn induce changes in soil polarization. Therefore, the medium polarization at low frequency may eventually reflect the root uptake. But this mechanism of measuring crop roots based on soil polarization is not verified.

Compared with the available literature at low frequency, information on the feasibility of using EC at high frequency to measure roots is quite limited. Basak and Wahid (2022) may be the first to report that the medium impedance at high frequency (5-100 kHz) is significantly negatively correlated with the root size (weight and length) of carrot with a root burying experiment. However, the mechanism and application of using medium polarization at high frequency to measure crop fine roots remains to be discovered. In a non-uniform system such as root and soil system, we know that root size, root uptake function and soil electrical parameters largely determine the system polarization. We can assume that when the effect of root absorption on soil electrical parameters is minimal (e.g. when fresh roots are buried in the soil), root size should correlate with the system polarization (here refers to soil capacitance, EC_soil_). However, root absorption decreases EC_soil_ in natural soil growth conditions and the absorption of water and nutrients by the roots relates to the size of the living root. Therefore, the effect of root polarization and absorption function on EC_soil_ are interacted with each other.

To date, it is unclear how root cell polarization and root absorption determine EC_soil_ at different frequency. If EC_soil_ can estimate root size, is it feasible for detecting root depth distribution under field conditions? Therefore, the aims of this study are to: (1) determine whether EC_soil_ can estimate root size; (2) explore the mechanism regulating EC_soil_ polarization at different measurement AC frequency; (3) determine whether EC_soil_ can measure the root depth distribution in the field.

## 2. Materials and methods

We conducted three experiments, namely root additions to aqueous solution or soil which we will refer to as “root immersing/burying”, soil pot and field experiments (Table 1). Field topsoil (0-20 cm) was collected at Luancheng Agro-Eco-Experimental Station of the Chinese Academy of Sciences (37° 53’ N, 114° 41’ E, altitude: 50.1 m) located in Shijiazhuang, China and then sieved through a 2.5 mm sieve to remove large plant litter for root burying and pot experiments. The soil texture is silty loam (Haplic luvisol according to FAO-WRB; IUSS Working Group 2015) and other detailed physicochemical properties can be found in Gu *et al*. (2021). The roots for the root immersing and root burying experiments were obtained from the field at Luancheng Agro-Eco-Experimental Station. The EC_soil_ of all experiments was measured at an output voltage of 1 V in parallel mode.

**Table 1.**
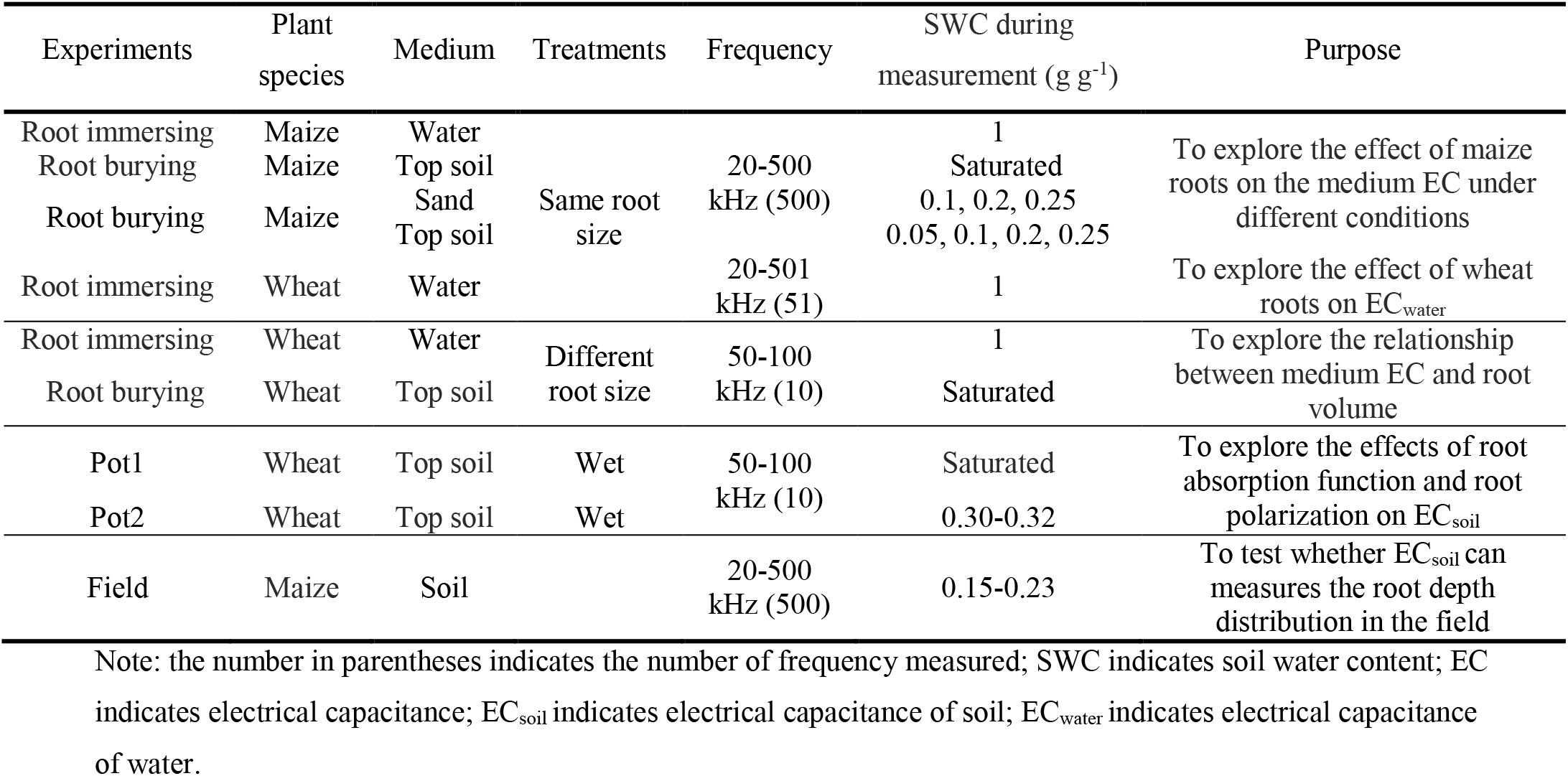
The experimental summary

### 2.1 Root immersing and burying experiments

To explore the potential mechanism underlying the relationship between EC and root properties, we conducted two series of root immersing and burying experiments. First, we used root immersing and burying to examine the effects of root additions on medium EC (Table 1). Then, we explored the correlation between EC and root volume.

Field grown maize (*Zea mays* L.) roots (15 days after sowing) were excavated with a shovel and washed with tap water gently. Fresh roots (60 g) were evenly separated into 6 portions and were put each into a plastic box (length in 12 cm, width in 8.6 cm and height in 5 cm) filled with tap water. Another six plots filled with tap water without roots were treated as blank control. The water EC (EC_water_) was measured using a LCR-meter (TH2826 with an accuracy of 0.1%, Tonghui Electronics Co., LTD, Changzhou, China) at 500 different frequencies linearly spaced from 20 Hz to 500 kHz with two stainless steel plates (10 cm long, 8 cm wide and 1 mm thick) spaced 11.8 cm apart used as electrodes (Figure 1a). The medium was changed from tap water to sieved soil at saturated water content (0.32 g g^-1^), and the EC_soil_ was measured again following the same protocol as EC_water_.

**Figure 1.**
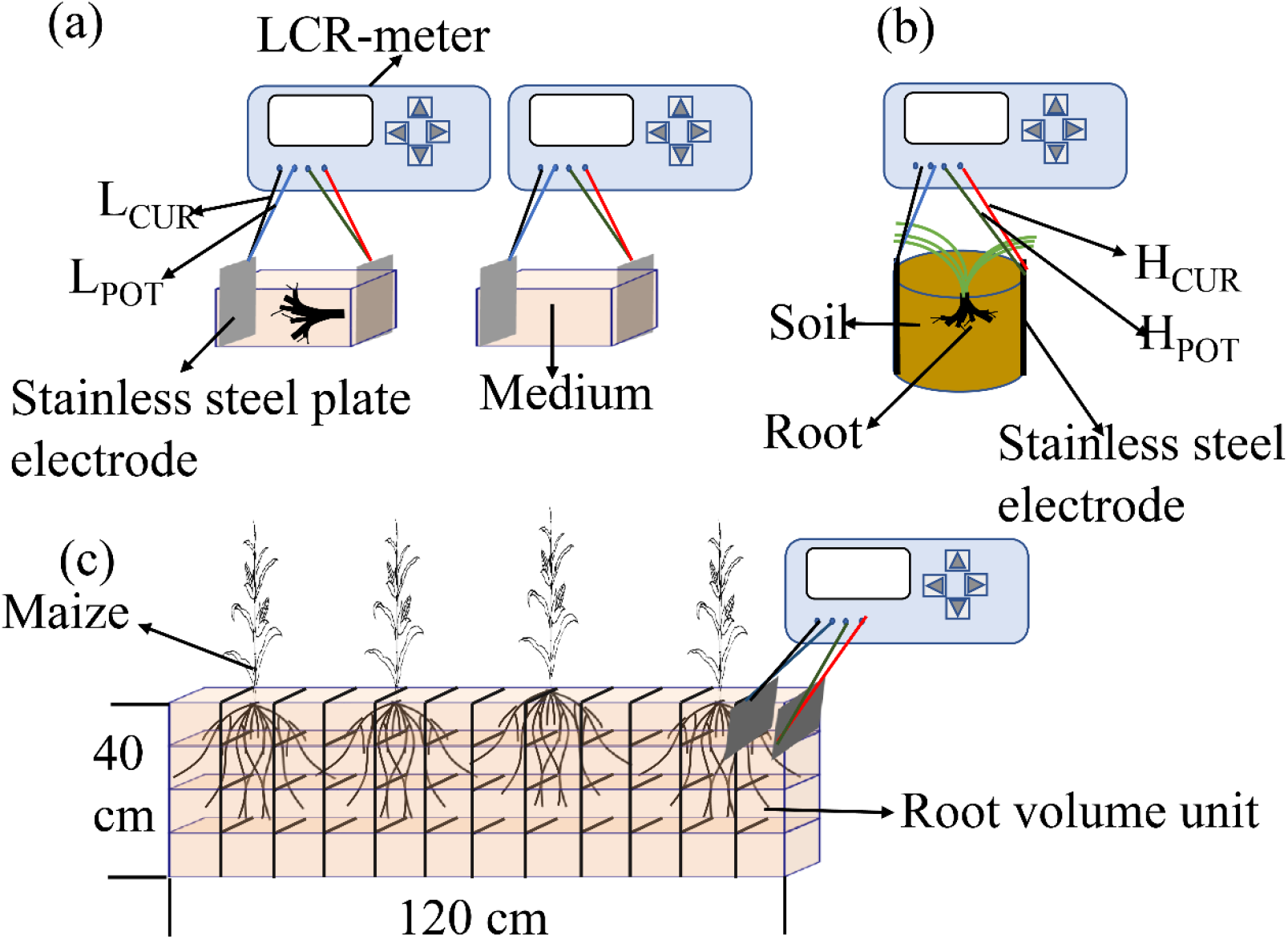
Schematic diagram of EC_soil_ measurements for different experiments; a indicates root immersing and burying experiments; b indicates soil pot experiments; c indicates field test experiment; L_CUR_ and H_CUR_ represent the low and high potential ends of the current output, respectively. L_POT_ and H_POT_ represent the low and high potential ends of the detection voltage, respectively; The measured soil sample volume is 10×10×8 cm^3^ in c.

Root burying experiment using maize roots (Table 1, 37 days after sowing) was conducted to explore the effect of root additions on EC_soil_ under different soil textures and water additions. We prepared sand with three different soil water contents (SWC) (0.1 g g^-1^, 0.2 g g^-1^ and 0.25 g g^-1^) and field sieved soil with four different SWCs (0.05 g g^-1^, 0.1 g g^-1^, 0.2 g g^-1^ and 0.25 g g^-1^) using dry sand and dry soil, respectively. Using same plastic pots above, 20 g of fresh roots were buried in the sand and field soil; each soil texture/water content combination was replicated six times. Similarly, another six replications of each soil texture/water content combination were created without root additions to serve as a control. The soil was then compacted to a bulk density of 1.4 g cm^-3^ and EC_soil_ was measured as described in the seedling maize root burying experiment.

We obtained winter wheat (*Triticum aestivum* L) roots (Table 1) from plants at the grain-filling stage from six different water treatments, as follows: no irrigation (W0), one irrigation application at the jointing stage (W1), two irrigations at the jointing and heading stages (W2), three irrigations at the overwintering, jointing, heading and filling stages (W3), four irrigation at the overwintering, jointing, heading and filling stages (W4), five irrigations at the overwintering, jointing, booting, heading and filling stages (W5). Four fresh root samples (10 g each) were weighed for each water treatment. The 24 root samples were placed in different plots in the plastic boxes and 500 ml of water was added to each plot. Meanwhile, another four plots filled with tap water without roots were treated as a control. The EC_water_ at 51 different frequencies from 20 Hz to 501.02 kHz in steps of approximately 10 kHz was measured using TH2826 LCR-meter.

Addition of wheat roots at the jointing stage were collected in the field to test the correlation between EC_water_ and root traits. The roots were cleaned as before and placed into 28 plots each with different quantities of fresh root mass. Then 500 ml of tap water was added to measure EC_water_. Four plots containing only water were used as blank control. We used the LCR-meter (TH2817B+ with an accuracy of 0.1%, Tonghui Electronics Co., LTD, Changzhou, China) to measure EC_water_ at 50 Hz, 60 Hz, 100 Hz, 120 Hz, 1 kHz, 10 kHz, 20 kHz, 40 kHz, 50 kHz and 100 kHz.

The wheat roots at the heading stage were collected again in the field to verify the relationship of EC_soil_ with root traits. After rinsing the roots, 12 groups of roots with different fresh mass were placed in plastic boxes as described above but filled the box with sieved soil and saturated with water. Again, four plots with bare soil were set up as blank control. EC_soil_ was measured similarly as EC_water_.

### 2.2 Soil pot experiments

Two soil pot experiments were conducted using wheat as a model crops. In the first soil pot (Pot 1) experiment, wheat cultivars with different growth characteristics named: Jimai 38, Jimai 22, Jimai 47 and Shixin633 were used to create variety of root quantities in the pots, 56 pots for each cultivar. Plants were propagated by placing one seed in a 20 ml pot, filled with 18 g of soil that was kept adequately watered. All of the pots were placed into a plastic tray (35 cm in long, 29 cm in wide) and watered every two days until the soil was saturated. There were 31, 50, 52 and 43 seedlings of Jimai 38, Jimai 22, Jinmai 47 and Shixin633, respectively.

For the second soil pot (Pot 2) experiment, the pot and soil used were the same as the first soil pot experiment, and the initial SWC was 0.11 g g^-1^. Six replicates of eight wheat cultivars: Gaoyou 2018, Nongda 212, Zhongmai 175, Shiyou 20, Jimai 22 Jimai 38, Jinmai 47 and Shixin633, were grown under wet soil moisture (0.32 g g^-1^) conditions. The cylindrical pots (16 cm in height, 14 cm in diameter) were filled with 2 kg of sieved soil and compacted to a density of 1.1 g cm^-3^ with an initial SWC of 0.10 g g^-1^. After filling the pot with dry sieved soil, 400 g of water was added to the soil to ensure a good soil moisture for seed germination. Six seeds were planted in each pot and the seedlings were thinned to three individuals after one week of germination. Another six pots without plants received the same water supply as a control. One week after emergence, plants were thinned to four in 32 pots, to two plants in eight pots and to one in eight pots.

### 2.3 EC measurement for the soil pot experiments

The TH2817B+ LCR-meter was used to measure EC_soil_ for both soil pot experiments (Figure 1b). For the Pot 1 experiment, the pot was saturated with water before the EC measurements. The stainless steel rods with a diameter of 3 mm and a length of 4 cm were used as electrodes to measure EC_soil_. For Pot 2 experiment, EC_soil_ and soil electrical resistance were measured at 20 days, 26 days, 31 days, 35 days, 38 days, 41 days, 44 days after sowing, using stainless steel rods with a diameter of 3 mm and a length of 18 cm as electrodes inserted 12 cm deep into the soil and spaced 13 cm apart. SWC was also measured at the same time as the finally EC_soil_ measurement.

### 2.4 Root traits measurement for the soil pot experiments

After the electrical parameters were measured, both of the pots were harvested. The soil was washed through a stainless steel sieve with a 0.425 mm mesh to collect the roots, they were then stored in a −20 °C freezer. Root samples were further cleaned in the laboratory by manually removing debris and scanned (Epson Expression 12000XL, USA) at a resolution of 600 dpi. Root parameters such as root length, root surface area and root volume were analyzed by WinRhizo software (ver. 2019a, Reagent Instruments Inc., Quebec, Canada). Then the roots were oven dried at 75 °C until constant weight. Root volume, biomass and length were highly correlated with each other, though root volume may be the primary basis for the EC hypothesis as indicated from the theory of the permittivity of the non-uniform system (Maxwell, 1891; Wagner, 1914). Therefore, root volume will be mainly used to represent root size hereafter.

### 2.5 Soil nutrient and ion measurements for the second pot experiment

We assumed EC_soil_ at low frequency sensed root uptake function as discussed in the introduction (generally, nutrients are taken up by plants in the form of ions). To verify this assumption, soil nutrient and ion concentrations were measured in the Pot 2 experiment. During the root harvest of Pot 2 experiment, aapproximately100 g of soil from each pot was collected and air-dried indoors. The soil total nitrogen and total phosphorus contents were measured by the Kjeldahl method and molybdenum-antimony resistance colorimetry, respectively. Soil total potassium content and soil soluble salt (SO4^2-^, Ca^2+^, Mg^2+^, Na^+^, K^+^) was measured by inductively coupled plasma mass spectrometry (Vanhoe *et al*. 1995). Cl^-^ was measured by titration with AgNO_3_. Electrical conductivity was measured using a handheld conductivity meter (Spectrum Technologies, Inc., USA).

### 2.6 Field test

Finally, to test whether EC_soil_ measures the root depth distribution in the field (Figure 1c), we dug a trench (120 cm in length in horizontal direction, 40 cm in depth and 60 cm in width) in a maize field at the heading stage. We used a TH2826 LCR-meter to measure EC_soil_ at 500 frequencies (20-500 kHz) with stainless steel plates as electrodes (10 cm length, 8 cm width and 1 mm thick). The electrodes were inserted 8 cm into the undisturbed soil and spaced 10 cm apart. The same paired electrodes were used to do all the measurements. There were 12 locations measured in the horizontal direction and 4 depth intervals (0-10, 10-20, 20-30 and 30-40 cm) in the vertical direction (the volume of soil sample is 10×10×8 cm^3^), a total of 48 data points. After the measurement, a knife was used to cut and remove sections of soil (10 × 10 × 8 cm^3^) to collect the roots from the volume. Additional soil samples were collected from nearby the root sampling sites and SWC was determined by the drying method (48 h at 105 °C). 24 soil rings (100 cm^3^) were used to collect intact soil samples at the four soil depth levels with six replications for each to determine soil bulk density (SBD). Soil volumetric water content (SVWC) was calculated by equation (1):

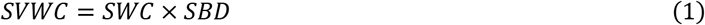

After completing the EC measurements, root samples were processed as the soil pot experiment and root traits were determined as well.

### 2.7 Statistical analysis

R (version 4.0.4) was used for statistics and plotting (R Core Team 2021). Before variance analysis, the data were tested for normality and homogeneity of variance and lm() functions were used for linear fitting. Least significant difference test (LSD_0.05_) was used for multiple comparisons. The t-test was used to compare whether the means were different and whether the correlation coefficient was zero. The change in EC (EC change rate) was derived from the following formula (2):

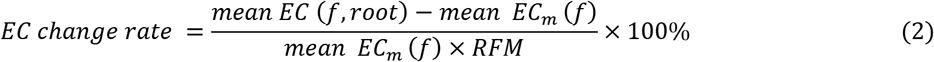

In the formula, mean EC (f, root) and mean EC_m_ (f) are mean EC with roots and without roots at f measurement frequency, respectively; RFM is root fresh mass.

## 3. Results

### 3.1 Effects of roots on medium EC at different frequency for the root immersing and burying experiments

To explore whether root volume affects medium EC, we first examined the effect of root existence on EC_soil_ and EC_water_ by the root immersing and burying experiments. The presence of maize and wheat roots did not significantly increase EC_soil_ or EC_water_ at low frequency (<1 kHz, Figure 2). As the measurement frequency increases, EC_soil_ and EC_water_ with roots buried in soil and immersed in water became significantly (*P* < 0.05, Figure 2) greater than those without roots, although soil and root properties also affected EC. Adding maize roots maximally increased EC_water_ by 44.9% while the maximum increase in EC_soil_ was 29.3% in the sand and 22.9% in field soil (Figure 2a). All these highest increments were observed at around 30 kHz. At the range frequency of 30 kHz −100 kHz, the increase of EC_water_ of wheat roots increased with more irrigation times during the whole growth period (Figure 2b).

**Figure 2.**
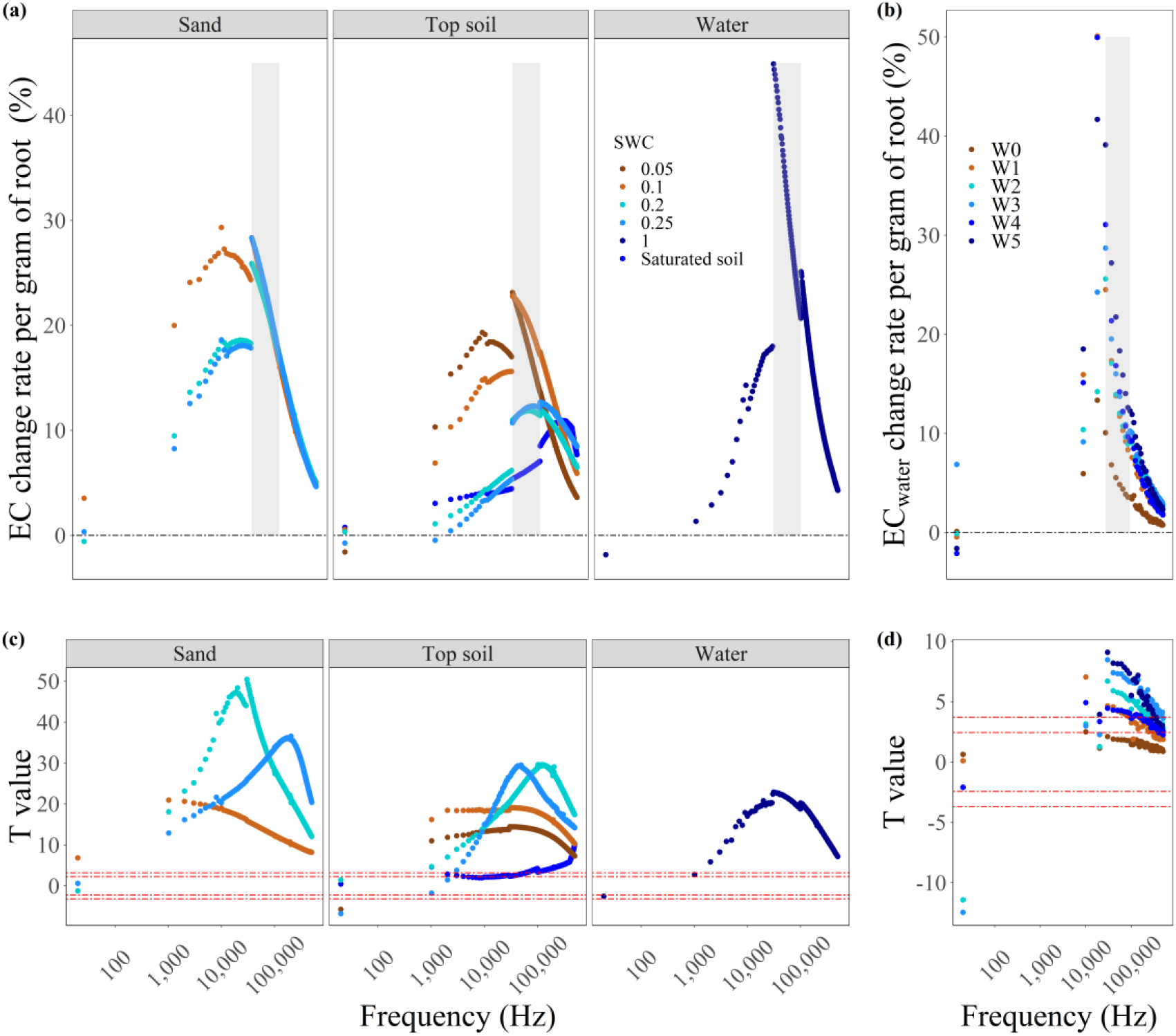
Effect of fresh roots on medium electrical capacitance (EC, a and b) and the corresponding two-tailed test results of mean values with and without roots in the medium (c, d); the shaded area represents the frequency ranges from 30 kHz to 100 kHz; W0, W1, W2, W3, W4 and W5 mean no watering, watering once, watering twice, watering three times, watering four times and watering five times during the whole growth period of wheat, respectively; The two red lines above 0 represent significant levels of 0.05 and 0.01, respectively. The shaded area represents the frequency ranges from 30 kHz to 100 kHz.

### 3.2 Relationships of root volume with EC_soil_ in the root immersing/burying and soil pot experiments

There was no significant correlation between root volume and EC_water_ or EC_soil_ at low frequency, however when the frequency was higher than 10 kHz, significant positive linear correlations between root volume and medium EC were observed in root immersing/burying experiments (Figure 3a and b). The correlation between medium EC and root volume was highest at 100 kHz (R=0.97 for both EC_water_ and EC_soil_, *P* < 0 .0001, Figure 3a and b).

**Figure 3.**
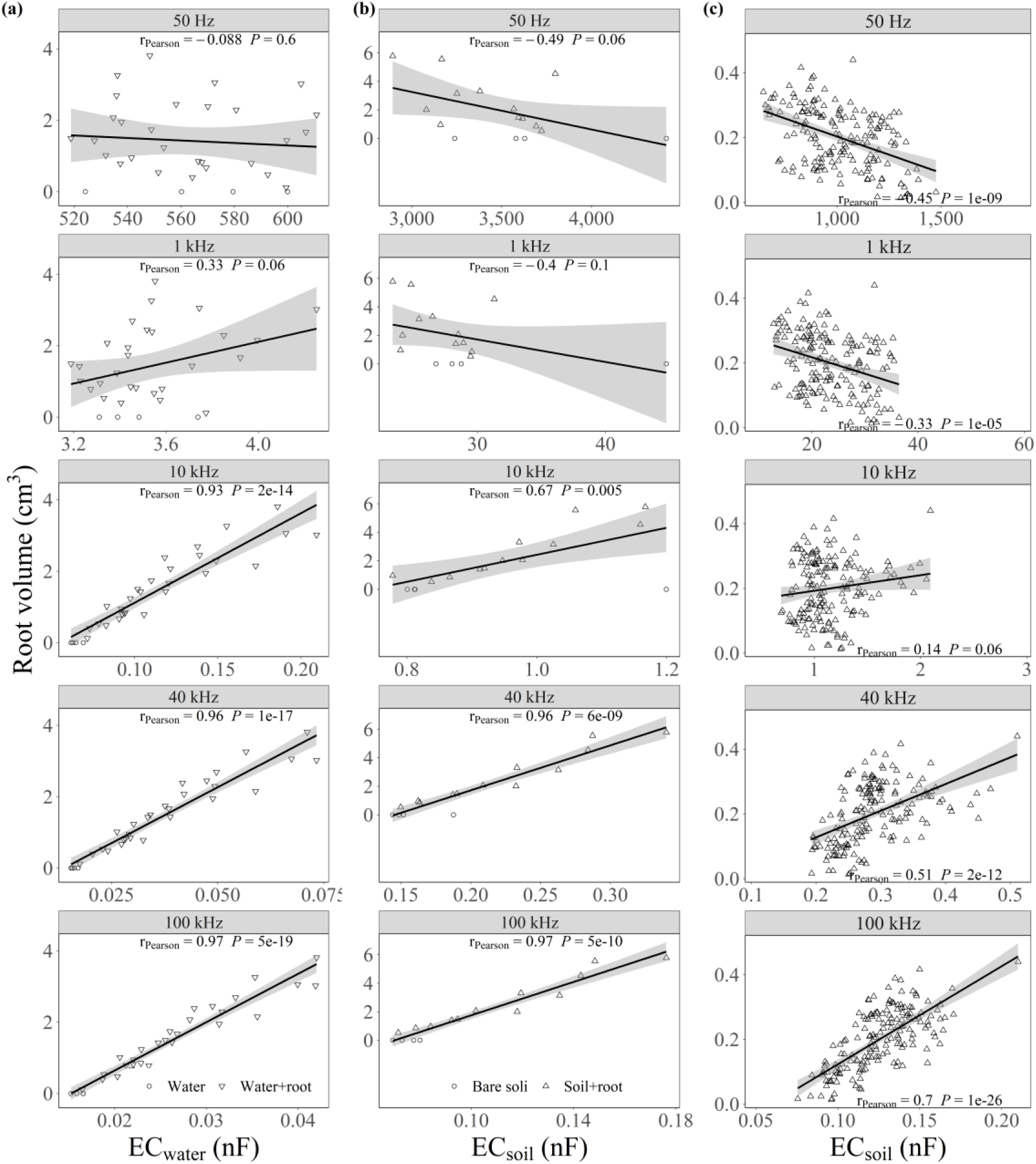
The relationships between root volume and EC_water_ (a, n=32) or EC_soil_ (b, n=16) at different frequency for the wheat root addition experiments and first soil pot experiment (c, n=176); EC_water_ and EC_soil_ indicate electrical capacitance of water and electrical capacitance of soil, respectively.

As a contrast, EC_soil_ at low frequency (<10 kHz) was significantly and negatively correlated with root volume (*P* < 0.05) in the soil pot experiments (Figure 3c and Figure 4a). When the measurement frequency was higher than 40 kHz, the correlations became positive and reached to the highest at 100 kHz (*P* < 0.01, Figure 3c and Figure 4a).

**Figure 4.**
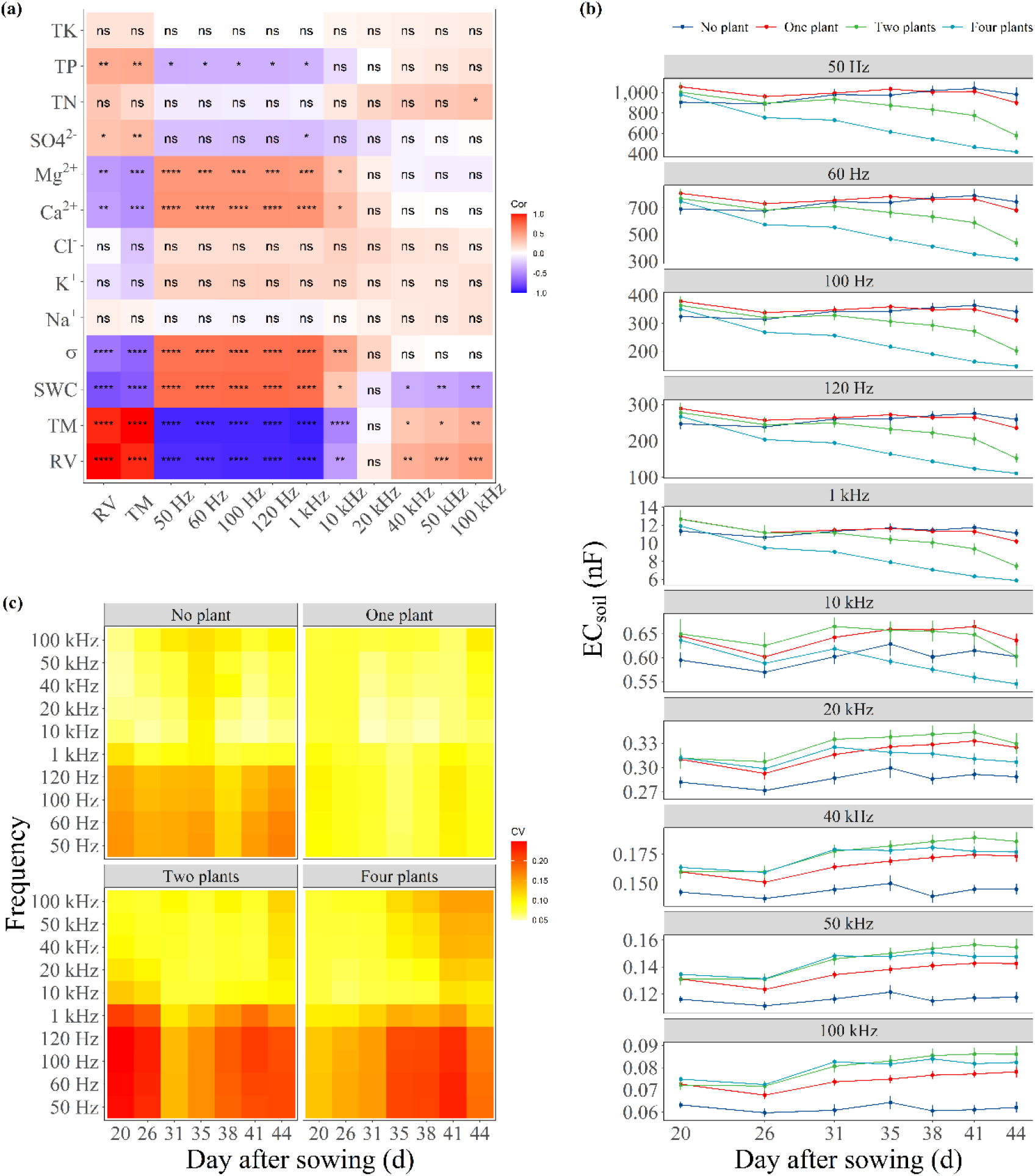
The correlation among soil electrical capacitance (EC_soil_) at different frequency (50 Hz, 60 Hz, 100 Hz, 120 Hz, 1 kHz, 10 kHz, 20 kHz, 40 kHz, 50 kHz, 100 kHz), root volume (RV), total biomass (TM, shoot dry mass plus root dry mass), and soil characters (SWC, σ, Na^+^, K^+^, Cl^-^, Ca^2+^, Mg^2+^, SO^42-^, TN, TP, TK) (a) and changes of EC_soil_ (b) and coefficient of variation (CV, c) among the replications with days after sowing in the second pot experiment; SWC, σ, Na^+^, K^+^, Cl^-^, Ca^2+^, Mg^2+^, SO^42-^, TN, TP, TK represent soil water content, saturated soil solution conductivity, soluble sodium ions, potassium ions, chloride ions, calcium ions, magnesium ions, sulfate ions, total nitrogen, total phosphorus and total potassium contents, respectively; Cor stands for linear correlation coefficient; ns, *, **, *** and **** represent the significant levels of insignificance, 0.05, 0.01, 0.001 and 0.0001, respectively; The vertical bar represents the standard error.

Furthermore, we found that there were significant negative correlations between root volume and SWC, soil electrical conductivity, soil soluble Ca^2+^ and soil Mg^2+^ content (*P* < 0.01) in the second soil pot experiment (Figure 4a). Resultingly, EC_soil_ at low frequency was positively correlated with SWC, soil electrical conductivity, soil soluble Ca^2+^and soil Mg^2+^ content proving that EC_soil_ at low frequency is a measurement of root uptake function. However, EC at high frequency was not significantly correlated with these soil characters except for SWC (Figure 4a). In addition, we also found that EC_soil_ with roots grown was smaller than EC_soil_ without roots grown when the frequency was lower than10 kHz, but it became larger than EC_soil_ without roots grown when the frequency was higher than 10 kHz (Figure 4b). The coefficients of variation EC_soil_ among the replications at high frequency (< 10 kHz) was smaller than that at low frequency (> 10 kHz, Figure 4c). These results enable EC_soil_ at high frequency to quantify root volume directly and accurately.

### 3.3 The field application

The EC method (EC_soil_) was evaluated under field conditions using maize (45 days old). There was a strong significant and positive correlation between root volume and EC_soil_ when the frequency was greater than 2 kHz (Figure 5a). The correlation coefficient between EC_soil_ and root volume were greater than 0.9 (*P* < 0.001), across frequencies ranging from 31 to 99 kHz with a maximum value of 0.92 at 71 kHz. When the frequency surpassed 100 kHz the correlation between root volume and EC_soil_ gradually decreased (Figure 5a). Using the best correlation at 71 kHz, we calculated the difference between measured and estimated root volume (Figure 5b and c). The mean error across all depths was15.0% with depth-specific means of 5.0%, 14.6%, 8.9% and 31.3% at 10 cm, 20 cm, 30 cm and 40 cm, respectively.

**Figure 5.**
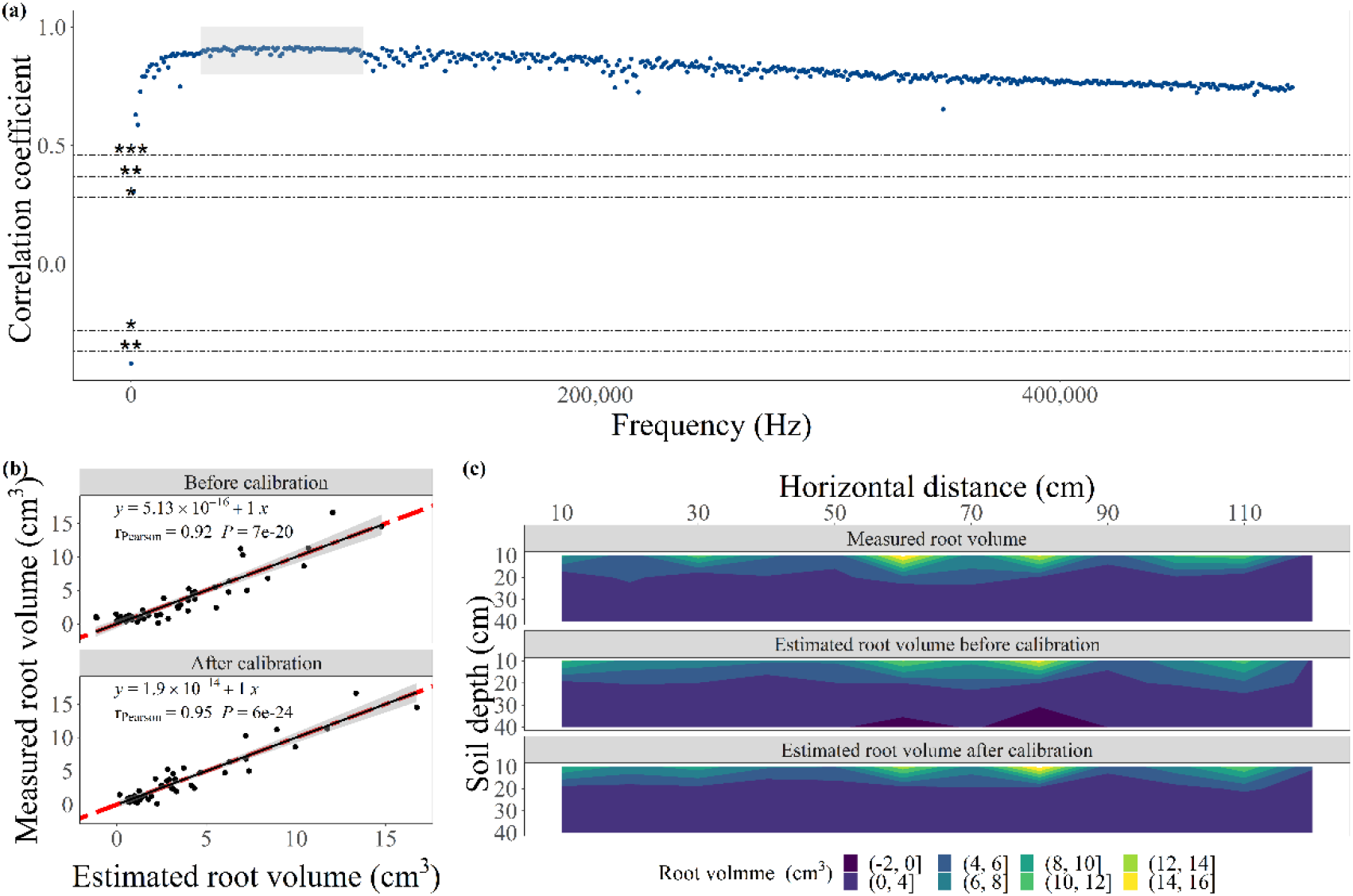
The correlation coefficients for the relations between root volume and soil electrical capacitance (a), the relationships between measured and estimated root volumes (b) and the comparison between measured and estimated root volume distribution (c) in the maize field experiment; *, ** and *** represent the significant levels of insignificance, 0.05, 0.01 and 0.001 respectively.

To improve the measurement accuracy it was necessary to consider and account for the effect of soil moisture on EC_soil_. Since there was a positive linear relationship between EC_soil_ and SVWC in the absence of roots (R^2^=0.99, *P* < 0.01, Table 2), we were able to calculate EC_soil_ (without roots) by EC_soil_ =0.248×SVWC+0.0127 and correct our field measurements. After SVWC calibration, the correlation coefficient between root volume and EC_soil_ increased from 0.92 to 0.95, and AIC and root mean square error declined, indicating that the calibration improved the correlation (Table 2). Using the calibration, root volume estimation based on EC_soil_ became more reliable (Figure 5c), and the depth specific errors for 0-10 cm, 10-20 cm, 20-30 cm and 30-40 cm soil depths were narrowed to 0.4%, 12.0%, 1% and 34%, respectively.

**Table 2.**
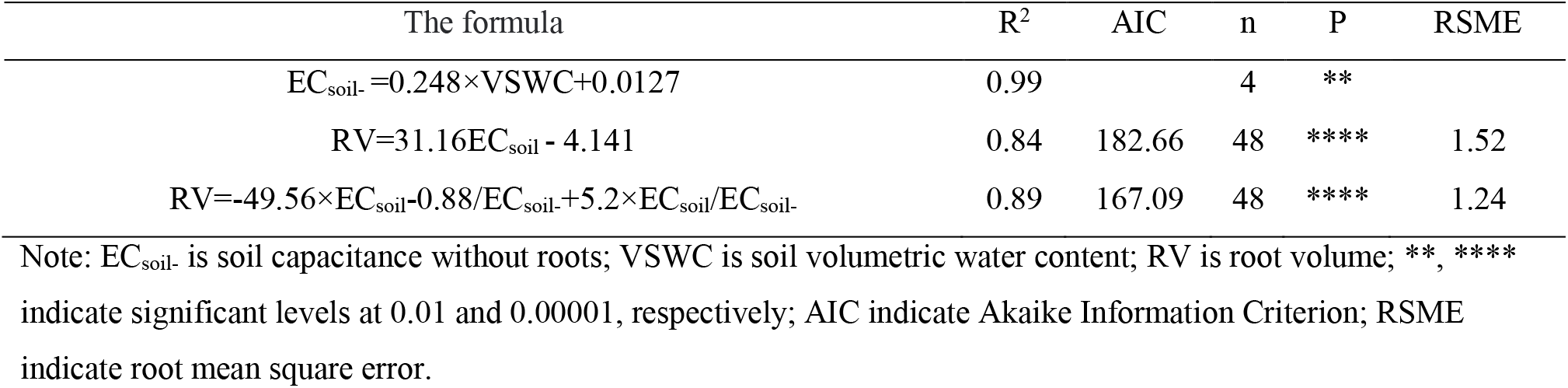
The relationships among soil capacitance, soil water content and root volume.

## 4. Discussion

### 4.1 EC correlates with root volume

Our results showed that the EC_soil_ at low (<1 kHz) or high frequency (30 kHz −100 kHz) can estimate root traits in the controlled and field conditions. Although roots did not directly increase EC_soil_ at low frequency (<1 kHz, Figure 2), the absorption of water and ions by roots can significantly reduce EC_soil_ (Figure 4a). Thus, root volume was negatively correlated with EC_soil_ both in the second soil pot and field experiments (Figure 3c, Figure 4a and Figure 5a). As the frequency is higher than 1 kHz, the presence of roots significantly increased EC_soil_ (Figure 2) indicating the dielectric constant of root is greater than the soil colloid, resulting in positive correlations between root volume and EC_soil_ for all the soil pot, root burying and field experiments (Figure 3, Figure 4a, and Figure 5a). Using the maximum influence of root presence on the medium EC as a benchmark, the optimal measuring frequency is between 30 kHz and 100 kHz for our sand and silty loam topsoil (Figure 2a). The results from the field experiment showed that EC_soil_ estimated surface root volume more accurately than deep root volume. Most of the roots reside and are distributed in the surface soil (0-20 cm); the paucity of roots deeper in the profile likely makes the increase of EC_soil_ smaller (Figure 5) while increasing the measurement error.

### 4.2 Mechanism of EC_soil_ measuring root volume

The root immersing and burying experiments showed that roots had not increase the medium EC at low frequency (< 1 kHz), which differs from previous reports (Mary *et al*., 2017; Weigand and Kemna, 2017; Weigand and Kemna, 2019; Tsukanov and Schwartz, 2021). Prior studies found that roots increased the polarization of solution or soil at low frequency, that is, increased the EC. This disparity may be explained by the use of nutrient solution to grow plants, which may have augmented root polarization (Ellis *et al*., 2013a) or simply the root systems used in prior experiments had a large root volume (> 10 cm^3^) (Mary *et al*., 2017). In this study, the volume fraction of root was small compared to the total volume of media, hence, the effects of root polarization on EC_soil_ were not significant (Figure 2). Another possible explanation is that soil porosity was increased in the process of root burying, which leads to the decrease of soil dielectric constant and thus EC_soil_ (Mary *et al*., 2017). Obviously, if plants grow in the medium such as in the soil pot experiment, the uptake of soil water and ions by roots will reduce the EC_soil_ (Ellis *et al*., 2013b), and the uptake amount is correlated with the root length and volume etc. Therefore, root traits can be indirectly measured based on EC_soil_ at low frequency.

Generally, the medium EC with roots is influenced by the interaction between the volume of roots and the medium EC without roots. As known, the electrical conductivity of root is greater than that of soil and solution (Weigand and Kemna, 2019; Basak and Wahid, 2022), so the current easily enters the root. When the frequency is lower than 100 kHz, the root cell membrane is insulated compared to the cell liquid phase, as a result, the root cell can be regarded as a conductive sphere covered by an insulating layer (Fricke, 1931; Mancuso, 2012) and root membranes act as charge-storing dielectrics of the capacitor (Dalton, 1995). When the frequency is greater than 100 kHz, the interfacial polarization is induced in the cell membrane and other membrane systems (Zhao, 2008). Additionally, the relaxation time of the soil is about 0.11 s (Mary *et al*., 2017) and its characteristic relaxation frequency is about 1.4 Hz. Therefore, the soil is not fully charged when frequency is higher than 1.4 Hz. With the increase of frequency, the soil becomes less charged, which is represented by the decrease of soil dielectric constant. However, when the frequency is less than 100 kHz, the cells will be fully charged (Zhao, 2008). Root epidermis and cell walls block the electrical charge movement and the charge accumulation leads to the interface polarization. Therefore, with the increase of frequency, the decrease of root permittivity is much smaller than that of soil permittivity. As a result, the root dielectric constant is much higher than that of the soil (Figure 2 and Figure 4b). This may be the reason that EC_soil_ at high frequency is linearly and positively correlated with root volume.

The polarization of root-soil system is not only affected by frequency, but also by root physiological state (Figure 2b) and soil environment. Under stress conditions, the increased permeability of the cell membrane leads to a decrease in the accumulation of charge on the cell membrane, which ultimately leads to a decrease in the permittivity of the root system (Zhao, 2008). It is also proved by our wheat roots immersing experiments (Figure 2b). SWC, soil ion content, soil bulk density, surface-charged colloid particles i.e. clay minerals and organic-mineral complexes can significantly affect EC_soil_ without roots (Li *et al*., 2005; Fares and Polyakov, 2006; Postic and Doussan, 2016, Cseresnyés *et al*., 2017), but we found that at high frequency, the reduction of EC_soil_ due to root absorption can be masked by root polarization (Figure 4b). This may partially explain why the correlations between EC_soil_ and electrical conductivity or ion concentration were not significant at high frequency in the second soil pot experiment (Figure 4a). In field studies it is important to consider the potential effects of soil heterogeneity, possibility pairing EC measures with SWC and EC measures without roots. Considering soil water into the prediction model, the correlation between EC_soil_ and root volume can be improved and the prediction error of root volume can be reduced in the maize field test (Figure 5b, Figure 5c and Table 2).

### 4.3 Future perspectives

As the field test provided us a reliable estimation of root depth distribution using EC method, it should be applicable to be extended to measure root 3D distribution and the dynamic changes in root growth, especially for the taproot systems which are difficult to assess with minirhizotron techniques. Compared with EC_root_, EC_soil_ does not need to consider current leakage in the root system which enables EC_soil_ to measure deep roots. Thus, we foresee that EC_soil_ opens a new door to develop nondestructive and noninvasive root phenotyping technique. However, to make the EC_soil_ more feasible, further systematic research on the influence of the optimal frequency, root physiological state, soil environment, and measurement method (electrode material, shape, distance etc) on the detecting accuracy is needed. For example, building the reversed model based on the same soil depths in the same soil type may help reduce the effect of soil bulk density, organic and SWC on EC_soil_ which can help improve the root detecting accuracy. Particularly, even if the equipment used in this study is not the most advanced one (< $3,900), it can still give us a satisfactory assessment of root traits. We believe with the development and application of new advanced equipment, the measuring accuracy should be improved further.

## 5. Conclusions

We proposed a new EC method to measure root volume based on root-soil system polarization. Whether under laboratory or field conditions, root polarization contributed greatly to soil polarization at high frequency (>10 kHz). Although the root polarization could be ignored at low frequency (<1 kHz), we speculate that the root uptake function affected soil polarization and may have some applications. Thus, root volume and other traits can be measured directly based on high frequency EC_soil_ and perhaps indirectly by EC_soil_ at low frequency. We believe the newly available equipment combined with the novel root predicting model will enable EC_soil_ to be a widely used root phenotyping technique in the future.

## Acknowledgements

The authors thank Hanbing Jiang, Shiming Duan, Weishuang Feng and Zhaoliang Mei for helping with field sampling and providing assistance in processing root samples.

## Author Contribution

H.G., X.L. designed research; H.G., B.L., X. Z., Y. L. and X.L. performed research; H.G. and X.L. analyzed data; all the authors wrote the paper.

## Data Availability

All study data are included in the main text and supplementary materials. Data can be downloaded from ScienceDB (https://www.scidb.cn).

## References

Basak R, Wahid KA. 2022. An In Situ Electrical Impedance Tomography Sensor System for Biomass Estimation of Tap Roots. Plants-Basel 11, 1713.

Cseresnyés I, Kabos S, Takács T, Végh KR, Vozáry E, Rajkai K. 2017. An improved formula for evaluating electrical capacitance using the dissipation factor. Plant and Soil 419, 237–256

Chloupek O. 1972. The relationship between electric capacitance and some other parameters of plant roots. Biologia Plant 14, 227–230

Cimpoiasu MO, Kuras O, Pridmore T, Mooney SJ. 2020. Potential of geoelectrical methods to monitor root zone processes and structure: A review. Geoderma 365, 114232

Dalton FN. 1995. In-situ root extent measurements by electrical capacitance methods. Plant and Soil 173, 157–165

Ehosioke S, Nguyen F, Rao S, Kremer T, Placencia-Gomez E, Huisman JA, Kemna A, Javaux M, Garre S. 2020. Sensing the electrical properties of roots: A review. Vadose Zone Journal 19, e20082

Ellis T, Murray W, Kavalieris L. 2013a. Electrical capacitance of bean (*Vicia faba*) root systems was related to tissue density-a test for the Dalton Model. Plant and Soil 366, 575–584

Ellis T, Murray W, Paul K, Kavalieris L, Brophy J, Williams C, Maass M. 2013b. Electrical capacitance as a rapid and non-invasive indicator of root length. Tree Physiology 33, 3–17

Fares A, Polyakov V. 2006. Advances in crop water management using capacitive water sensors. Advances in Agronomy 90, 43–77

Fricke H. 1931. The electric conductivity and capacity of disperse systems. Physics 1, 106–115

Gan Y, Liu L, Cutforth H, Wang X, Ford G. 2011. Vertical distribution profiles and temporal growth patterns of roots in selected oilseeds, pulses and spring wheat. Crop & Pasture Science 62, 457–466

Gu H, Liu L, Butnor JR, Sun H, Zhang X, Li C, Liu X. 2021. Electrical capacitance estimates crop root traits best under dry conditions-a case study in cotton (*Gossypium hirsutum* L.). Plant and Soil 467, 549–567

IUSS Working Group. 2015. International soil classification system for naming soils and creating legends for soil maps. World Soil Resources Reports 106, FAO, Rome

Karlova R, Boer D, Hayes S, Testerink C. 2021. Root plasticity under abiotic stress. Plant Physiology 187, 1057–1070

Kessouri P, Furman A, Huisman JA, Martin T, Mellage A, Ntarlagiannis D, Bücker M, Ehosioke S, Fernandez P, Flores-Orozco A et al. 2019. Induced polarization applied to biogeophysics: recent advances and future prospects. Near Surface Geophysics 17, 595–621

Li X, Lei T, Wang W, Xu Q, Zhao J. 2005. Capacitance sensors for measuring suspended sediment concentration. Catena 60, 227–237

Liao A, Zhou Q, Zhang Y. 2015. Application of 3D electrical capacitance tomography in probing anomalous blocks in water. Journal of Applied Geophysics 117, 91–103

Lynch JP. 2013. Steep, cheap and deep: an ideotype to optimize water and N acquisition by maize root systems. Annals of Botany 112: 347–357

Lynch JP. 2019. Root phenotypes for improved nutrient capture: an underexploited opportunity for global agriculture. New Phytologist 223, 548–564

Mancuso S. 2012. Measuring roots: An updated approach. Springer Science & Business Media, Springer-Verlag Berlin Heidelberg

Mary B, Abdulsamad F, Saracco G, Peyras L, Vennetier M, Meriaux P, Camerlynck C (2017) Improvement of coarse root detection using time and frequency induced polarization: from laboratory to field experiments. Plant and Soil 417: 243–259.

Maxwell JC. 1891. A Treatise on Electricity & Magnetism. Oxford UK: Clarendon Press

Peruzzo L, C. Chou W, Wu YX, Schmutz M, Mary B, Wagner FM, Petrov P, Newman G, Blancaflor EB, Liu X et al. 2020. Imaging of plant current pathways for non-invasive root Phenotyping using a newly developed electrical current source density approach. Plant and Soil 450, 567–584

Postic F, Doussan C. 2016. Benchmarking electrical methods for rapid estimation of root biomass. Plant Methods 12, 1–11

R Core Team. 2021. A language and environment for statistical computing. Vienna, Austria: R Foundation for Statistical computing. http://www.R-project.org/

Tsukanov K, Schwartz N. 2021. Modeling Plant Roots Spectral Induced Polarization Signature. Geophysical Research Letters 48, e2020GL090184

Tsukanov K, Schwartz N. 2020. Relationship between wheat root properties and its electrical signature using the spectral induced polarization method. Vadose Zone Journal 19, e20014

Urban J, Bequet R, Mainiero R. 2011. Assessing the applicability of the earth impedance method for *in situ* studies of tree root systems. Journal of Experimental Botany 62, 1857–1869

Vanhoe H, Saverwijns S, Parent M, Moens L, Dams R. 1995. Analytical characteristics of an inductively coupled plasma mass spectrometer coupled with a thermospray nebulization system. Journal of Analytical Atomic Spectrometry 10, 575–581

Wagner K. 1914. Explanation of the dielectric fatigue phenomenon on the basis of Maxwell’s concept. Arkiv für Electrotechnik 2, 371–387

Weigand M, Kemna A. 2017. Multi-frequency electrical impedance tomography as a non-invasive tool to characterize and monitor crop root systems. Biogeosciences 14, 921–939

Weigand M, Kemna A. 2019. Imaging and functional characterization of crop root systems using spectroscopic electrical impedance measurements. Plant and Soil 435, 201–224

Zhao K. 2008. Dielectric spectroscopy and its applications. Beijing China: Chemical Industry Press

